# Influence of Contact Map Topology on RNA Structure Prediction

**DOI:** 10.1101/2024.11.25.625138

**Authors:** Christian Faber, Utkarsh Upadhyay, Oskar Taubert, Alexander Schug

## Abstract

The available sequence data of RNA molecules have highly increased in the past years. Unfortunately, while computational power is still under exponential growth, the computer prediction quality from sequence to final structure is still inferior to the labour intensive experimental work. Although a reliable end-to-end procedure was already developed for proteins since Alphafold2, while its successor AlphaFold3 can also predict RNA, its confidence in particular for novel sequences and folds appear still limited. Another strategy entails two steps: (i) predicting potential contacts in the form of a contact maps from evolutionary data, and (ii) simulating the molecule with a physical force field while using the contact map as restraint. However, the quality of the structure prediction crucially depends on the quality of the contact map. Until now, only the proportion of true positive contacts was considered as a quality characteristic. We propose to also include the distribution of these contacts, and have done so in our recent studies. We observed that the clustering of contacts, as is common for many AI algorithms, has a negative impact on prediction quality. In contrast, a more distributed topology is beneficial. We have applied these findings from computer experiments to current algorithms and introduced a measure of distribution, the Gaussian score.

## 1 Introduction

According to the central dogma of molecular biology, RNA functions as an intermediate product in protein biosynthesis [1]. In the following years, however, more and more observations were made that RNA fulfils further functions. The influence of the structure of tRNA, for example, was already discussed in the 1980s [2], or the structure of the ribosomal RNA [3]. These RNA molecules, that do not code for a protein, are called non-coding RNA (ncRNA) [4]. Similar to proteins, the spatial structure of ncRNA is crucial for its ability to function successfully [5]. Therefore, the correct determination of the structure is of fundamental importance for the understanding of ncRNA, including evolutionary paths [6], virus genesis [7] and also the design of new drugs [8, 9]. The problem is that unlike sequencing, the experimental determination of the structure is a very labour-intensive task [10]. Therefore, a *in silico* approach that requires less effort would be a major advance and achievement.

Immense progress has been made in the field of protein structure prediction in the last decade. Due to the large amount of available data, it was possible to train large language models and thus achieve an end-to-end prediction that comes close to experimental quality. The pioneering implementation of google with *AlphaFold* [11] should be mentioned in this context. In the case of RNA structure prediction, such an assessment is rather problematic due to data limitations [12, 13]. To illustrate the difference in the data situation, we can look at the published PDB structures in the central database *rcsb*.*org* : For proteins we have 196, 400 structures and for RNA only 1, 890 (status: November, 2024). As the end-to-end machine learning approaches are not yet fully developed, are there other possibilities? Various physics based methods are available to determine the three-dimensional structure of RNA molecules. The most prominent representative for the structure folding, but also function determination, of molecules is the molecular dynamics (MD) simulation [14, 15, 16]. For example, it can be used to describe the full dynamics of RNA folding[17, 18]. So while molecular dynamics simulations are very accurate, when used in the context of structure prediction, they require very large amounts of computing time, even when introducing biasing constraints [19, 20]. For this reason, simplified models are often used in practice for structure prediction, which can determine the stationary folding more quickly using Monte Carlo algorithms. Similarly, much effort has been spend into judging the quality of RNA structure prediction[21]. We studied here how to achieve reliably good predictions in a realistic computing time. To achieve this, we use a workflow that has been successfully used in protein structure prediction since before *AlphaFold*. In the following, we will describe this process in a little more detail; for a better understanding, it is shown graphically in Figure 1.

**Figure 1.**
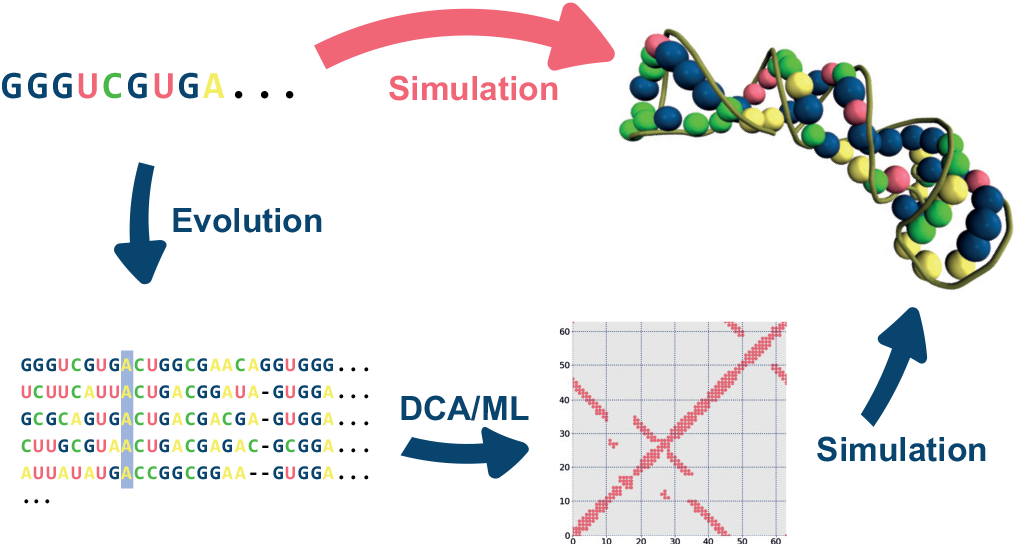
Schematic representation of the typical RNA structure prediction workflow. The upper route produces significantly worse results without the detour via contact map creation.

Two popular prediction software are the Monte Carlo-based SimRNA [22] and the fragment-based program Rosetta [23]. Both are capable of converting the initial sequence directly into a three-dimensional RNA structure. With them, the structure prediction quality can vary considerably with excellent predictions of low RMSD but also, e.g., only achieving an RMSD of ∼ 29^°^A for *3q3z*. Evolutionary data can support predictions by acting as helpful additional biasing constraints and considerably improve the quality of predictions [24, 25] by assuming that co-evolutionary mutational patterns are a results of spatial adjacency. To detect co-evolutionary signals, homologous sequences from other organisms are collected for the initial sequence 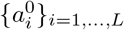 with 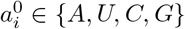 and organised as multiple sequence alignment (MSA) 𝒟 = {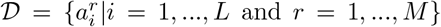 and *r* = 1, …, *M* } with 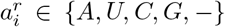, *M* the number of sequences. The co-evolution of different residues can be analysed with methods used in statistical physics, for instance mean field direct coupling analysis (DCA) [26, 27, 24], or with modern AI algorithms [28, 29, 30]. These methods give us a binary mapping CM : ℤ_*L*_ *×*ℤ_*L*_ → ℤ_2_, called contact map, which indicates whether two residues *i, j* are spatially adjacent (the nitrogen atoms are less or equal 9.5^°^A apart), i.e. form a contact. In simulations, these contacts can be used as restraints. The outcomes of these simulations are notably superior when evolutionary data is incorporated. Thus, the task is on the one hand to have a good data basis for the creation of MSAs and on the other hand to develop algorithms that provide as much information as possible from the MSA to the simulation software. In the past, the latter was often implemented using the direct coupling analysis method mentioned above, with positive predictive values (PPV) of around ∼ 50% being observed [31, 32, 33]. The co-evolutionary analysis can also be carried out using machine learning algorithms. The PPVs of these algorithms reach values of up to 80% [29, 28], which is a remarkable increase. Due to the high PPV, one would assume an improved structure prediction by the simulation. However, detailed investigations were unable to confirm this expectation [29]. In fact, the increased PPV through a convolutional neural network (CNN) brought almost no improvement in prediction quality compared to the DCA-generated contact map. This result could have two causes. Firstly, the prediction quality could have an upper limit due to the simulation software, i.e. a further improvement of the restraints is unnecessary. On the other hand, the contact maps generated by AI algorithms could be of poorer quality, despite a high PPV. The AI algorithms are trained to achieve the best possible PPV, but the PPV is a massive dimension reduction of the initial situation. The PPV calculates a value from the contact map of the experimental structure and the contact map of the evolutionary analysis, i.e. 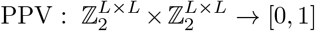. Machine learning methods, like convolutional neural networks [34], suggest that contacts may be predicted in clusters wherever possible, therefore the PPV of such contact maps is still high, but information about smaller contact areas is lost. In this paper we want to investigate whether this effect of contact clusters is really responsible for the reduced performance in structure prediction and to introduce a mathematical measure to prevent this effect. This measure can be integrated into the objective function of ML algorithms to generate more diverse topologies of contact maps. To address these questions, we first present the software used and our test set in Materials and Methods. Followed by explaining different methods to model various kinds of topological different contact maps for our testset and introduce the new measure to gauge the diversity. We also describe the design of the computer experiments and how the influence of false contacts was investigated. Finally, the different applications are presented. In the Results and Discussion section, the results of the computer experiments are discussed and the value of the new measure is shown directly in the applications. In the Conclusion, we summarise the most important findings again, but also address weaknesses and problems of the computer experiments carried out. Finally, we discuss possible continuations in modern AI algorithms.

## 2 Materials and Methods

In the following section we give an overview of our test setup and the different computer experiments we ran on the test set to gain insight into the simulation quality in relation to different contact map topologies.

### 2.1 Simulation Software and Test-Data

As simulation software, we use SimRNA [22], a software based on replica exchange Monte Carlo (REMC), with the standard configuration and ten replicas per simulation. We use the representative of the largest cluster as the final structure. The contacts are added as restraints in the energy function. A description of the penalty term can be found in the SI.

The presented computer experiments were performed with a test-subset 𝔇 of RNA families composed in reference [33]. We had to sort out the structures that had a faulty PDB structure and thus made a comparison with our simulation results impossible. This effectively gave us 56 RNA families representing non-coding RNA molecules ranging in length from 41 to 496 residues. In the application section, we use a different test set 𝔇_Val_. Furthermore, three additional data points from the test set, designated as 𝔇_Val_, are excluded from the evaluation process due to the lack of good MSAs for these data points (we refere to the reduced dataset as 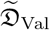). Furthermore, the AI algorithm used was trained with parts of our current test set 𝔇. This ensures that we can make a statement that is as independent as possible. In this section we have also adapted our contact definition for the same reasons. An overview of all families,representing molecules, their size, the effective family size *M*_eff_ and contact definitions is provided in the SI.

### 2.2 Distribution of Contacts

The results of CoCoNet [29] suggest that the distribution of contacts has a direct influence on the prediction of the spatial structure of the RNA molecule with *L* nucleotides. In order to investigate this influence, three distinct contact distributions are generated for each molecule within the test set 𝔇. The individual distributions are constructed by generating the contact map from the experimentally determined three-dimensional structure and selecting a subset of *L/*2 contacts from it. These *L/*2 contacts are selected in the following ways: [label=()]

1. Clustered
2. Randomly
3. Gaussian weighted

This gives us three different contact map topologies for each molecule. How well the three different topologies improve the simulation and whether the simulation benefits at all from this information will be discussed in the Results section.

The **clustered** contact maps are an extreme example of the way a convolutional neural network, for example, operates contact prediction. The probability of finding a contact is increased if there is already a contact in the immediate vicinity. We select our contacts from the complete contact map in such a way that we only take contacts from the largest contact cluster. If these are already all selected, we take contacts from the second largest cluster and so on. An example of such a selection is drawn in figure 3 (a).

**Figure 2.**
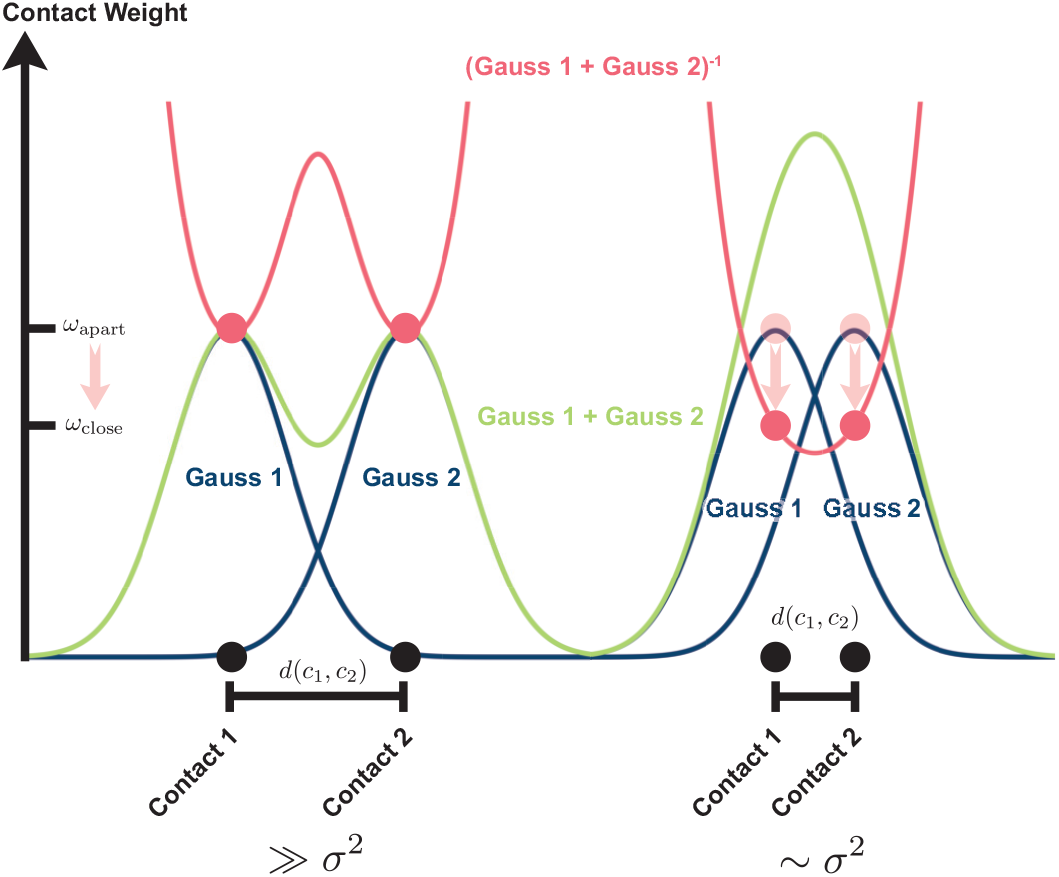
One dimensional schematic of the Gaussian contact weighing. On the left are two contacts with a distance *d* much larger than *σ*^2^. On the right are the contacts closer together and the overlap of the individual Gauss distributions (blue) is bigger. After summation (green) and inverting (red) the weights *ω*_close_ decrease compared to *ω*_apart_ for the contacts further away.

**Figure 3.**
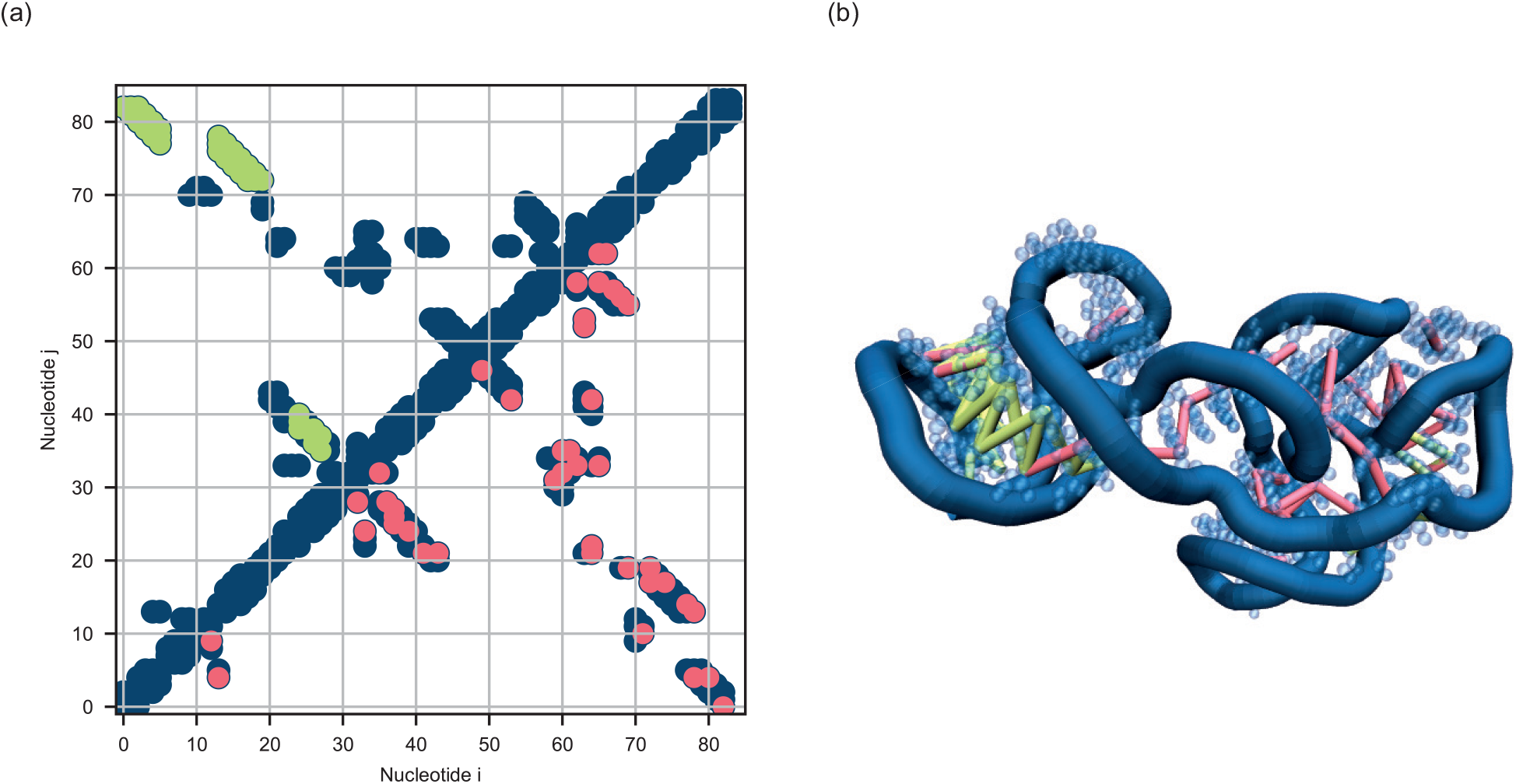
**(a)** Contact map of the *cobalamin riboswitch aptamer domain* (ID: *4frg*). In darkblue is the native contact map as derived from crystal structure. In green a possible choice of restraints forming clusters and in red randomly distributed contacts. **(b)** The three-dimensional structure of the molecule *4frg* from the test set 𝔇, determined experimentally. The coloured bonds represent the contacts belonging to (a).

The **randomly** distributed contacts represent the opposite extreme case. In this distribution, the individual restraints are randomly selected from the entire contact map. This random selection is very difficult to incorporate into existing algorithms. What is needed is a measure that determines the degree of randomness.

We introduced this measure with a **Gaussian** weighting of the individual contacts. Instead of valuing each contact equally, contacts that are close to each other are devalued. A one dimensional schematic representation of this can be seen in Figure 2. Two scenarios are depicted there: on the one hand, two contacts in the one-dimensional contact map that are further apart and, on the other hand, those that are close together. The influence of each contact on the neighbouring contact is modelled with a Gaussian function (blue curve) and the addition of all influences of the other contacts gives the weighting of the current contact (green curve). To devalue the individual weights instead of enhancing them, this weighting is inverted (red curve). Here, the value at the point of contact can be read directly from the curve and the closer two contacts are to each other, the lower the value (hereafter referred to as the Gauss score). Generalised to two dimensions, we obtain the value ν_*i*_ for the addition of all Gaussian functions for each contact *i* with position *r*_*i*_ on the contact map with all contacts *C*:

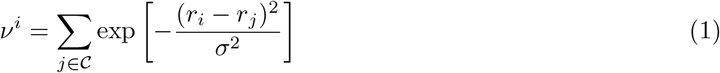

Each contact, which is surrounded by other contacts, has now a higher ν compared to isolated contacts. To devalue the contacts we introduce the inverse 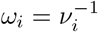 as weights. The total sum of weights is now reduced for a clustered contact selection and is at its maximum for widely dispersed contacts. The latter is closer to a random selection of contacts, so you can incorporate

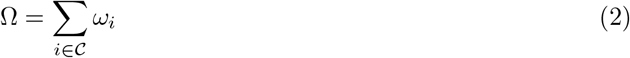

as an additional optimisation parameter in machine learning algorithms for the contact diversity. This parameter is a measure of the topology of the contact map. It can be used to evaluate existing algorithms, as we do in section 3.3, or to develop new algorithms that use it as an additional optimisation parameter. This is also the advantage over randomly distributed maps, which, although they are maximally spreaded, have no quantitative measure that can be optimised in calculations.

### 2.3 Influence of False Contacts

In reality, it is impossible to use only true positive contacts, since even the best current machine learning methods only achieve a PPV of around 80% [28]. In the next step we introduce false contacts to our contact map. We therefore use randomly choosen contacts 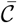 from the set of residue pairs which are not part of the native contacts. The remaining contacts, designated as set *C*, are generated through a Gauss optimisation process, derived from the native contacts. The positive predicted value (PPV) decreases from 1 to 1 − *λ*, where 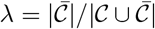 is the fraction of false contacts in our selection (error rate). With the selection 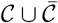 we can simulate each molecule d from our testset and calculate the 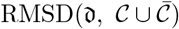. To give a short overview of the quality we introduced the beneficial fraction

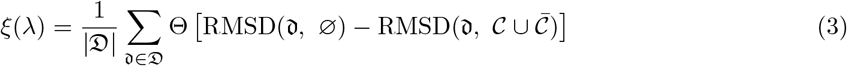

which shows the fraction of molecules for a given *λ* with a lower RMSD with given restraints compared to the same molecule without any restraints. In order to stay as close as possible to reality, we have decided on Gaussian-optimised contacts for the true positives and random for the false positives. This should best represent the influence of the false positives.

### 2.4 Applications

In order to analyse the individual influences and results on the structure prediction and the Gaussian score in practice, we looked at and compared two state of the art methods for contact map creation. As a benchmark, we start with the Direct Coupling algorithm, which has a lower PPV but should have a diverse contact map topology. We used the implementation pyDCA with the mean field approximation (mfDCA). As a representative of a sparse learning algorithm (Convolutional Neural Network) we take CoCoNet, which, as already mentioned in the introduction, has extremely good PPV values but not much gain in structure prediction [33]. The third method uses the advanced AI algorithm Barnacle, which has been specially trained for contact card prediction. Barnacle uses a transformer model to create a prediction that is as accurate as possible. For all three methods, we compare the precision and the Gauss value and take a closer look at contact maps. In order to reproduce the previous results from the literature [33, 28], we also switch to a different contact definition.

We count two residues *i, j* as contact iff there is a pair of heavy atoms that are less than 10 ^°^A apart.

## 3 Results and Discussion

### 3.1 Distribution of Contacts

We investigated the initial question of whether the influence of the distribution of contacts has an impact on the prediction quality by simulating different distributions of contacts. We investigated the three different distributions clustered, randomly and Gauss optimised and as presented in section Materials and Methods, and entered the results for all individual molecules into a common overview (see fig. 4). For comparison, the simulation without restraints is also presented, and it is clear that almost without exception the addition of restraints improves the prediction quality. The distribution with the clustered contacts gives only a slight improvement in the prediction. However, the Gaussian optimised distribution shows a significant improvement in the RMSD. Only two molecules (*3r4f* and *4qln*) do not benefit from the Gaussian contact distribution. However, the quality of these molecules is either already in a very high range (*4qln*) or both clustered and Gaussian contacts are worse than without restraints (*3r4f*), which points to a more fundamental problem. For all other molecules, the Gaussian-optimised distributions are in the range of randomly selected contacts. We can draw two conclusions from these results. Firstly, our hypothesis that a more diverse contact map contributes significantly to prediction quality is correct.

**Figure 4.**
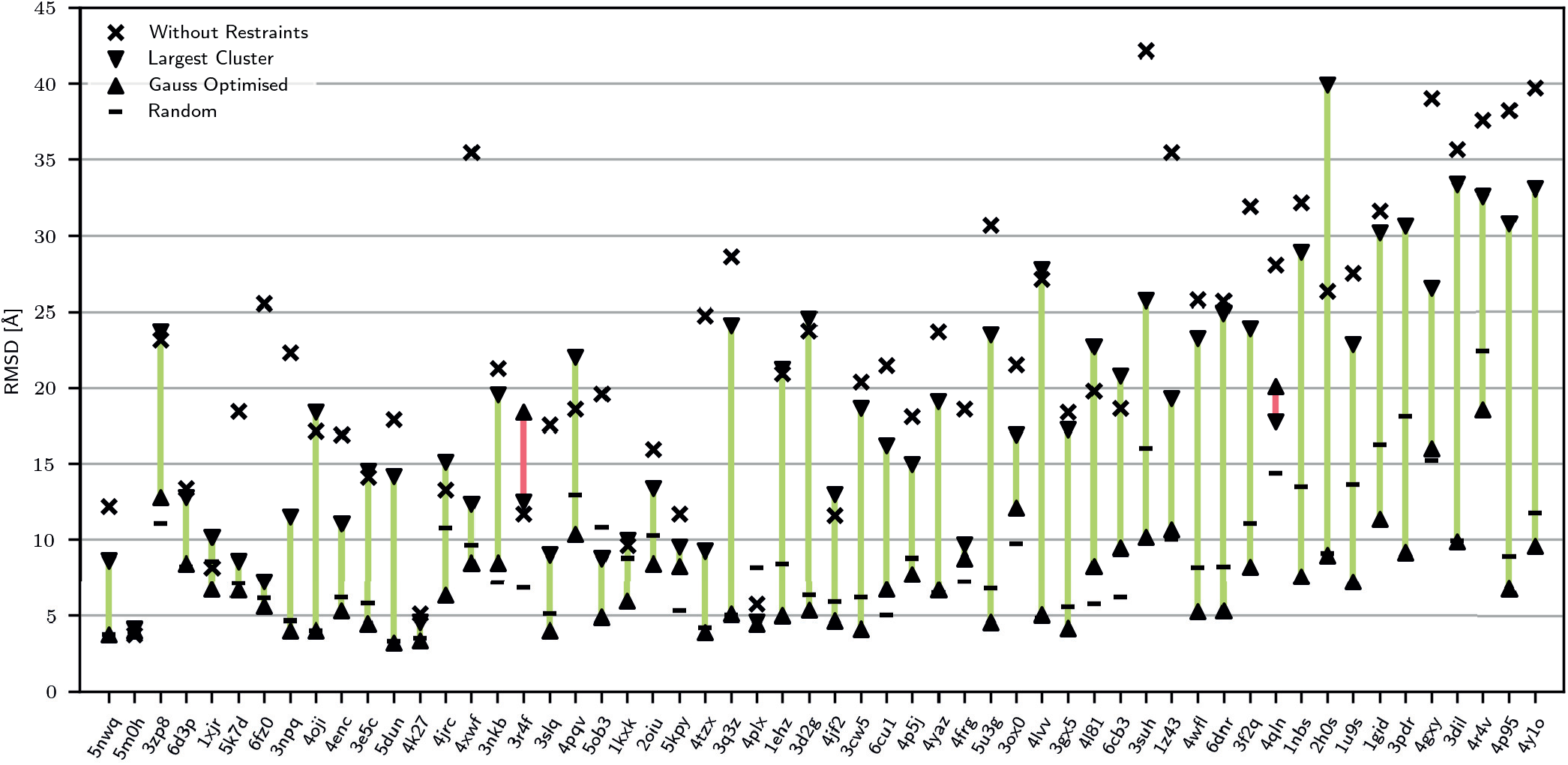
Complete visualisation of the prediction quality for all molecules for different distributions of contacts. The molecules are sorted by size and increase in size from left to right. The ids from *Protein Data Bank* [35] are plotted on the x-axis. All distributions have been derived from the native contacts from experimentally determined structures. We used different methods for this: Random, Clustered and Gaussian optimised. Simulations without any restraints are also shown for comparison.

Secondly, our Gaussian weighting Ω (see eq. 2) is a useful property of a contact map to evaluate the diversity or randomness of the same. Before we turn to the results of the specific applications, we want to analyse the influence of false contacts.

### 3.2 Influence of False Contacts

Assuming we include false contacts in our contact maps, the false contacts should also lead to a worse prediction. In Figure 5 (a), we used the beneficial fraction from equation 3 to analyse the PPV at which it is better not to specify any restraints for the simulation of an unknown molecule. The diagram demonstrates that even for a high error rate *λ* of 0.5, 70% of the molecules still benefit from the restraints. If we go into more detail and divide our test-set 𝔇 into larger molecules and smaller molecules using a threshold. We use the randomly selected values of 60, 75, 100 and 150 residues as the threshold. All molecules smaller than or equal to this value show a worse beneficial fraction than the larger ones in the analysis, with the exception of the very small molecules at very low error rates. However, this may also be due to the statistical significance of the very small sample. We see that the larger molecules in particular benefit from the additional restraints. For instance, the eight largest molecules profit exclusively from the inclusion of restraints, even though half of all contacts are false contacts. This admittedly very superficial analysis provides neither explanations nor does it say anything about specific molecules. But it shows that even with the most unfavourable choice of false contacts, their inclusion statistically leads to a better prediction. It is therefore highly advisable not to use a pure simulation, especially for larger molecules, but to perform an evolutionary analysis beforehand. Ideally, this evolutionary analysis should have as diverse a contact distribution as possible, as shown in the previous section. An detailed list of all individual molecules can be found in the SI.

**Figure 5.**
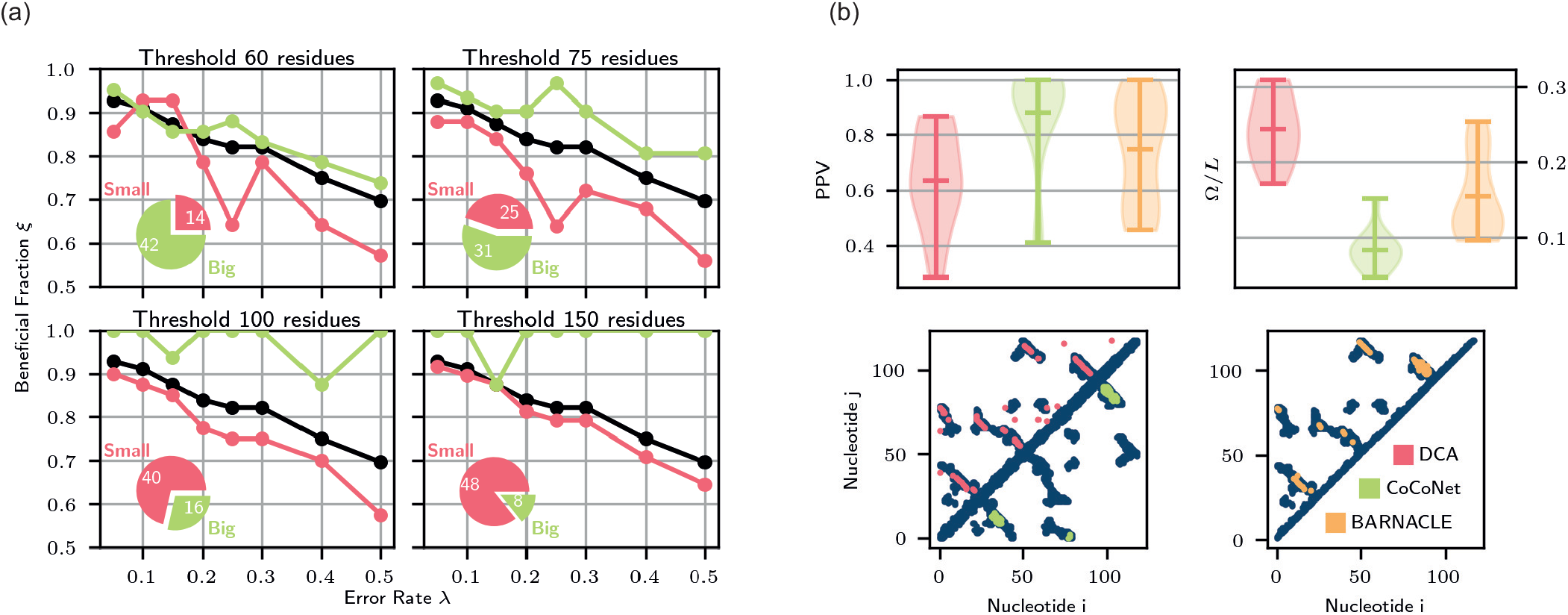
**(a)** Illustrations of the beneficial fraction. In black is the whole test-set for comparison, in green the bigger molecules above the specific threshold and in red the smaller portion with less or equal number of residues. The pie chart is a visual representation of the ratio and number of molecules for the individual subsets. **(b)** Study of the individual quantities (PPV and Ω*/L*) of the predictions on the test set 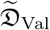. The bottom line shows the exemplary contact maps for the molecule *6ues* for all three prediction methods. The interpretation of the values is given in the main text. A listing of the validation set and the analysis of all individual molecules is given in the SI.

### 3.3 Applications

We have seen in the previous section that the inclusion of evolutionary data is always advantageous, even if the false contacts are maximally unfavourably located. The first section has already shown that the topology of the contact map has a decisive influence on the quality of the structure prediction. Finally, let us look again at familiar ML algorithms CoCoNet [33] and Barnacle [28] and try to explain the lack of structure prediction quality by including the topology of the contact map. First of all, let’s get a rough overview of the complete validation set. Figure 5 (b) shows the precision as an established quality value and the Gaussian value introduced by us, normalised to the size of the molecule, for quantifying the topology. As assumed in the introduction, it is already apparent at first glance that the ML algorithms, especially CoCoNet, have a more clustered topology compared to the DCA algorithm. But you can also see a difference between the two ML algorithms. The shallow algorithm CoCoNet leads to very poor Gauss values. In contrast, the algorithm Barnacle, which is based on a transformer architecture, provides a significantly higher Gauss value. If we look at a contact map in detail using the *6dues* molecule as in figure 5 (b), this distinction is immediately apparent. DCA generates many widespread contacts, even in the intermediate spaces, and thus achieves a low PPV, but in return has a large Gauss score due to the broad distribution. CoCoNet, in contrast, only recognises contacts in three separate clusters, whereas Barnacle provides a prediction that is almost perfectly distributed. This also emphasises the importance of including the contact map topology in the design of AI algorithms for structure prediction. Really good results and unnecessary detours can only be achieved with the inclusion of a good topology and not with pure PPV orientated training. In the already very successful algorithms for protein structure prediction, consideration of the topology may already be priced in by the large amounts of training data, but these amounts do not exist for RNA and the use of such knowledge is elementary for a good prediction.

## 4 Conclusion

We investigated whether the topology of contact maps, which are output by modern AI algorithms for RNA structure prediction, has an influence on the prediction quality. To answer this question, we created artificial contact maps and used a large test set to show that the topology has an important influence. To quantify the topology, we also introduced a new measure, the Gaussian score. In the last section, we applied the measures PPV and Gauss to a smaller validation set. At this point, however, it should also be said that the validation set in particular is of poor quality. In some cases, the underlying MSA consisted of only a few entries. Therefore, the absolute values in particular should be treated with caution. The individual contact maps are also shown in the SI, and in some of them you can see a completely inaccurate prediction of contacts. Another limitation of our analysis is the dependence on the simulation software SimRNA. However, the aim was not to achieve absolute numbers and the best possible prediction, but rather to compare different starting conditions as independently as possible. This is solved very well with SimRNA, since the simulation follows a physical force field, and not learned motives, as with trained AI algorithms. For RNA, we have the big problem of limited data sets for training LLMs, so an approach like AlphaFold is not yet feasible. Although AlphaFold 3 [30] has the option of RNA structure prediction, initial tests show that its quality does not match the prediction quality of proteins [36].

It is therefore of great importance to have clues for a good structure prediction in advance, in order to compensate for the lack of training data. The incorporation of topology is one such clue, which is of particular importance for architectures such as convolutional neural networks. In the future design of AI algorithms for structure prediction, on the one hand, such insights can be integrated and provide direction for the network structure. On the other hand, the Gauss score can also be used directly as a candidate term in the loss function. This allows the algorithm to directly train an optimised topology for ideally matched contact maps for structure prediction.

Lastly, DCA identifies co-evolutionary signals within RNA families, even to the extend of judging fitness landscapes [37]. The Gauss score provides a quantitative measure of the extent to which evolutionary constraints act delocalized across the contact map. Thus, it can be employed to assess whether evolutionary processes preferentially stabilize localized clusters of contacts or exert their influence in a more dispersed manner across distinct regions of the contact map.

## 5 Data Availability

The data underlying this article are available in ConMapTop at https://github.com/KIT-MBS/conmaptop.

## 7 Author contributions statement

C.F. performed all simulations, all authors analysed the results, C.F. and A.S. wrote and reviewed the manuscript.

## 8 Supplementary Data

Supplementary Informations are available.

## 9 Acknowledgments

Simulations were performed with computing resources granted by RWTH Aachen University under project rwth1287. The authors also gratefully acknowledge the Gauss Centre for Supercomputing e.V. (www.gauss-centre.eu) for funding this project by providing computing time through the John von Neumann Institute for Computing (NIC) on the GCS Supercomputer JUWELS at Jülich Supercomputing Centre (JSC) [38]. AS thank also recognizes NIC for funding this project. For parts of the paper, DeepL, a translation AI, was used.

