## Supplementary Information for "Influence of Contact Map Topology on RNA Structure Prediction"

### Influence of Contact Map Topology on RNA Structure Prediction - Supplementary Information

November 2024

#### 1 Software and Data

All software scripts, data and instructions how to use the code are stored in the following Git repository: https://github.com/KIT-MBS/conmaptop

#### 2 Contact Definition

In the main part of our work, we used the following contact definition:

> *Two nucleotides $i$ and $j$ are considered to be in contact if their nitrogen atoms are within $9.5\AA$ of each other.*

In the Applications section, however, we have adapted our contact definition to that of the original publication of the various methods. It reads:

> *Two nucleotides $i$ and $j$ are considered to be in contact if the nearest pair of atoms $(a, b)$, where $a$ is a heavy atom of nucleotide $i$ and $b$ a heavy atom of nucleotide $j$ are less than $10\AA$ apart.*

#### 3 Penalty Function SimRNA

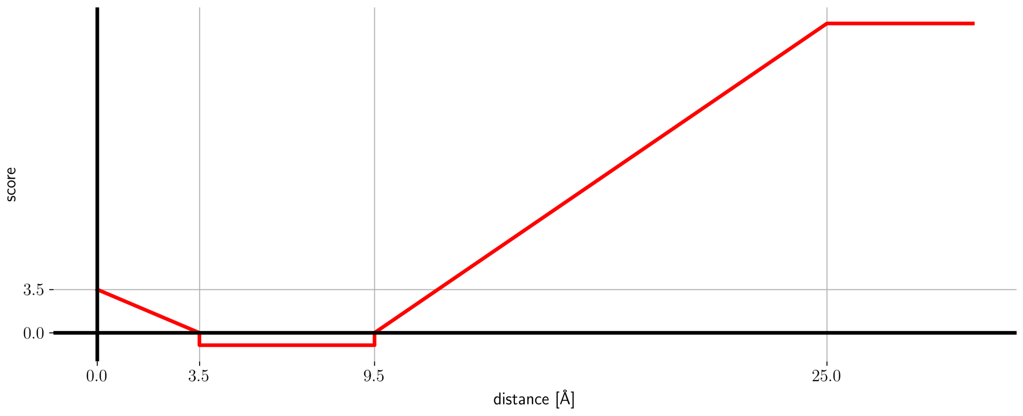


Figure 1: Each potential contact is weighted with this function in SimRNA.

### 4 Datasets

#### 4.1 Test Set $\mathfrak{D}$

Table 1: Dataset for the main part of the work.

| **PDB** | **Family** | $L$ | $M_{\text{eff}}$ |
|---|---|---|---|
| 1ehz | RF00005_1 | 76 | 461.0 |
| 1gid | RF00028 | 159 | 799.3 |
| 1kxk | RF00029 | 70 | 51.0 |
| 1nbs | RF00011 | 120 | 157.6 |
| 1u9s | RF00010 | 155 | 1285.3 |
| 1xjr | RF00164 | 47 | 3.3 |
| 1z43 | RF00169 | 101 | 281.5 |
| 2h0s | RF00234 | 123 | 603.7 |
| 2oiu | RF03017 | 71 | 2.0 |
| 3cw5 | RF00005_2 | 77 | 705.0 |
| 3d2g | RF00059 | 77 | 1241.9 |
| 3dil | RF00168 | 173 | 1455.5 |
| 3e5c | RF01767 | 53 | 13.6 |
| 3f2q | RF00050 | 108 | 104.8 |
| 3gx5 | RF00162 | 94 | 322.1 |
| 3nkb | RF02682 | 64 | 41.0 |
| 3npq | RF01057 | 51 | 10.5 |
| 3ox0 | RF00504 | 87 | 847.6 |
| 3pdr | RF00380 | 161 | 119.6 |
| 3q3z | RF01786 | 75 | 372.1 |
| 3r4f | RF00044 | 66 | 2.0 |
| 3slq | RF01510 | 67 | 6.6 |
| 3suh | RF01831_2 | 101 | 198.5 |
| 3zp8 | RF00163 | 43 | 59.0 |
| 4enc | RF01734 | 52 | 221.1 |
| 4frg | RF01689 | 84 | 62.8 |
| 4gxy | RF00174 | 162 | 6987.2 |
| 4jf2 | RF01054 | 77 | 14.6 |
| 4jrc | RF02447 | 57 | 6.8 |
| 4k27 | RF02695 | 55 | 32.0 |
| 4l81 | RF01725 | 96 | 211.0 |
| 4lvv | RF01831_1 | 89 | 187.7 |
| 4oji | RF02684 | 52 | 4.7 |
| 4p5j | RF00233 | 84 | 16.2 |
| 4p95 | RF01807 | 189 | 3.2 |
| 4plx | RF02266 | 76 | 2.0 |
| 4pqv | RF01415 | 68 | 4.9 |
| 4qln | RF00379 | 117 | 998.9 |
| 4r4v | RF02927 | 186 | 2.0 |
| 4tzx | RF00167 | 71 | 458.9 |
| 4wfl | RF01854 | 106 | 177.0 |
| 4xwf | RF01750 | 64 | 130.7 |
| 4y1o | RF02001_2 | 258 | 248.3 |
| 4yaz | RF01051 | 84 | 583.4 |
| 5dun | RF00921 | 54 | 2.0 |
| 5k7d | RF02679 | 47 | 32.4 |
| 5kpy | RF01982 | 71 | 2.0 |
| 5m0h | RF00606 | 42 | 2.0 |
| 5nwq | RF01763 | 41 | 5.1 |
| 5ob3 | RF01300 | 69 | 2.0 |
| 5u3g | RF00442_1 | 85 | 33.9 |
| 6cb3 | RF00080 | 99 | 409.5 |
| 6cu1 | RF02553 | 80 | 91.4 |
| 6d3p | RF02888 | 45 | 1.0 |
| 6dnr | RF00442_2 | 107 | 137.8 |
| 6fz0 | RF01826 | 48 | 2.8 |

#### 4.2 Validation Set $\mathfrak{D}_{\text{Val}}$

Table 2: Dataset for the application section.

| **PDB** | **Family** | $L$ | $M_{\text{eff}}$ |
|---|---|---|---|
| 1c2x | RF00001 | 120 | 5884.9 |
| 5di2 | RF00008 | 48 | 156.4 |
| 1l9a | RF00017 | 128 | 929.3 |
| 1p6v | RF00023 | 68 | 278.6 |
| 5zal | RF00027 | 73 | 34.8 |
| 6ol3 | RF00102 | 112 | 29.7 |
| 2mf0 | RF00166 | 72 | 36.8 |
| 4gma | RF00174 | 210 | 7636.0 |
| 2ke6 | RF00207 | 48 | 1.0 |
| 4c4q | RF00209 | 233 | 4.0 |
| 2nbx | RF00210 | 108 | 26.7 |
| 1s9s | RF00374 | 101 | 9.6 |
| 1z2j | RF00480 | 45 | 3.3 |
| 2krl | RF00500 | 102 | 2.0 |
| 6ues | RF00634 | 119 | 191.6 |
| 2lc8 | RF01073 | 63 | 24.0 |
| 2n1q | RF01381 | 155 | 2.0 |
| 6qn3 | RF01704 | 50 | 42.5 |
| 6hag | RF01727 | 43 | 14.4 |
| 5ddp | RF01739 | 61 | 37.6 |
| 3ndb | RF01857 | 136 | 136.5 |

### 5 Contact Maps

All the contact maps from the application section:

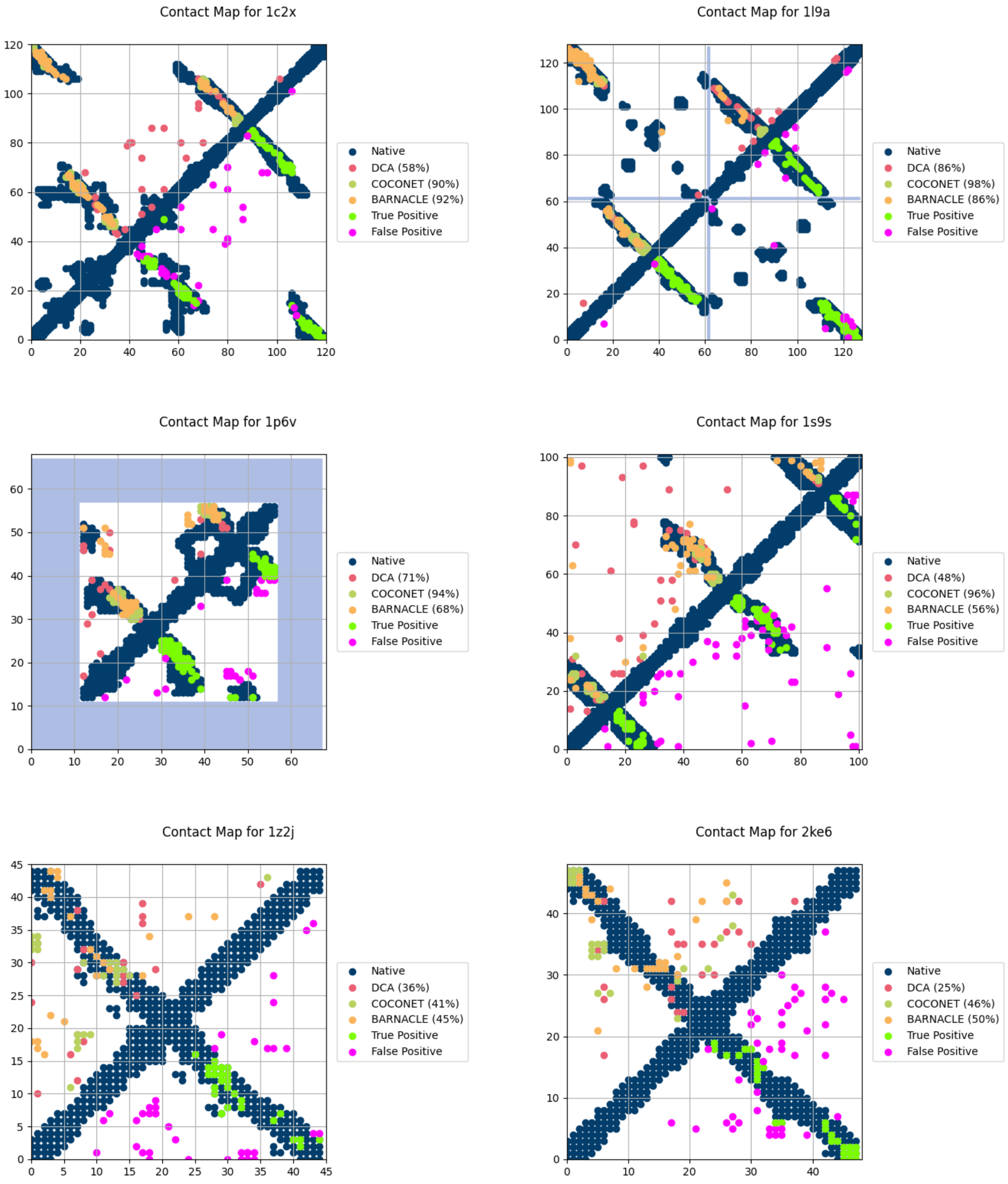


Figure 2: Contact Maps

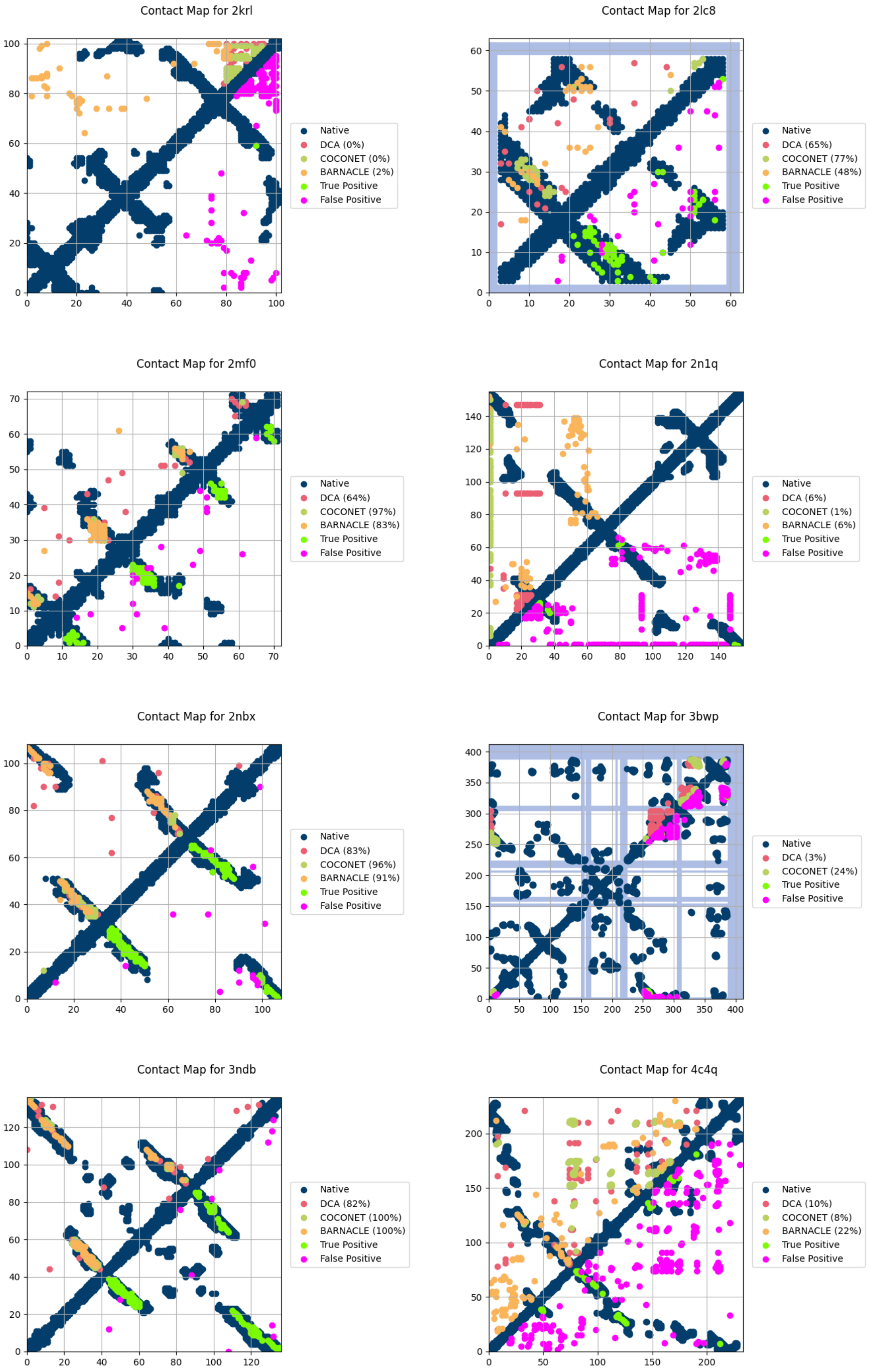


Figure 3: Contact Maps

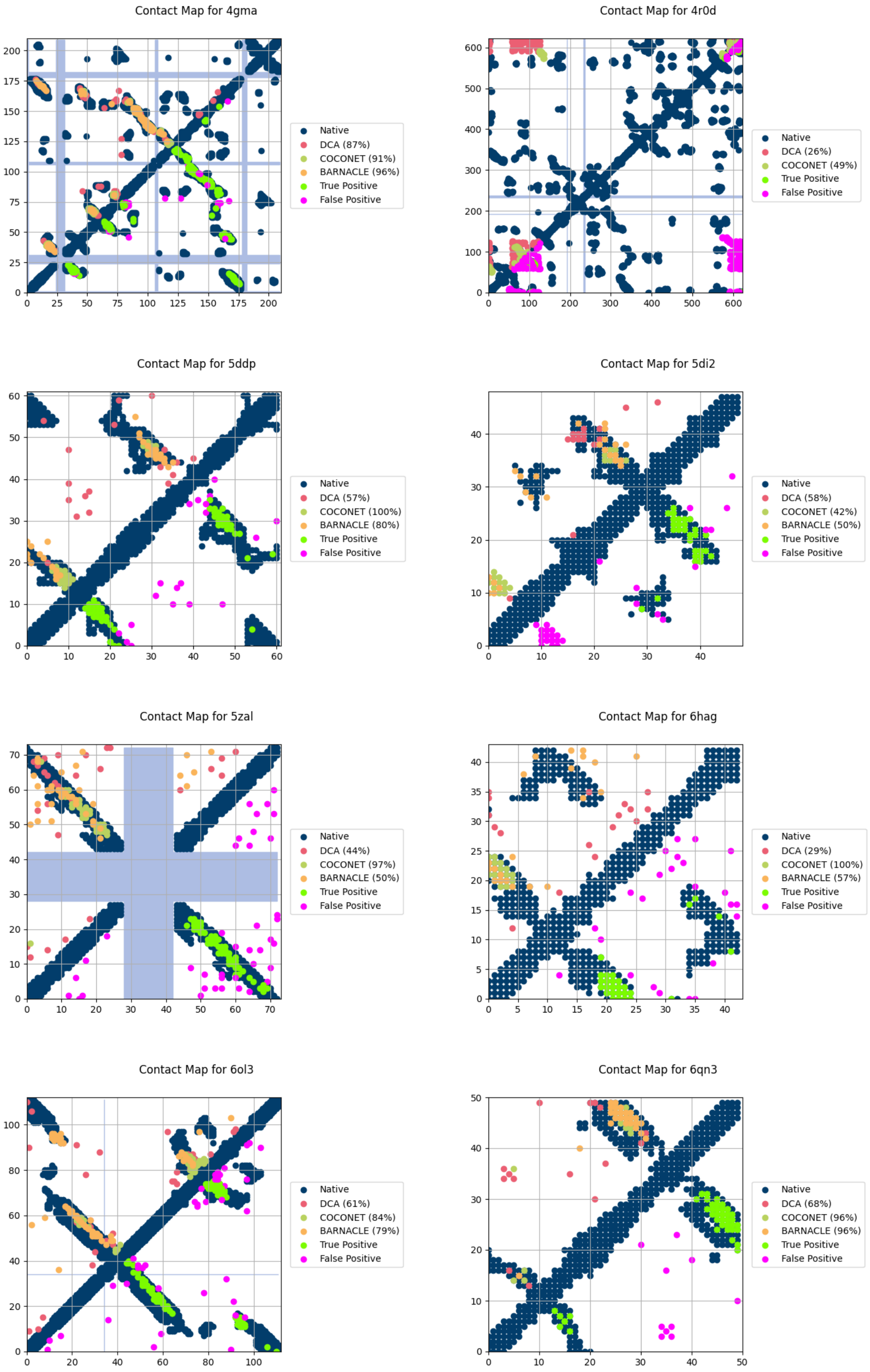


Figure 4: Contact Maps

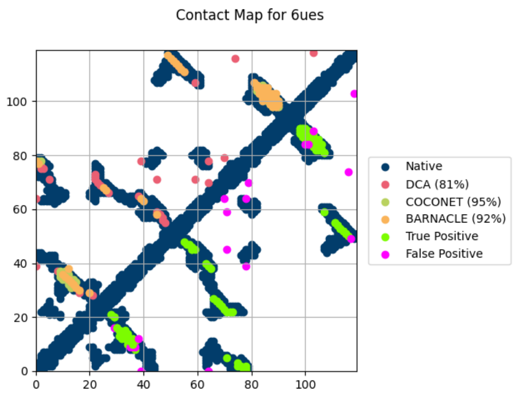


Figure 5: Contact Maps
